# Acoustic and postural displays in a miniature and transparent teleost fish, *Danionella dracula*

**DOI:** 10.1101/2021.11.10.468077

**Authors:** Rose L. Tatarsky, Zilin Guo, Sarah C. Campbell, Helena Kim, Wenxuan Fang, Jonathan T. Perelmuter, Eric R. Schuppe, Hudson K. Reeve, Andrew H. Bass

## Abstract

Acoustic communication is widespread across vertebrates, including among fishes. We report robust acoustic displays during aggressive interactions for a laboratory colony of *Danionella dracula*, a relatively recently discovered miniature and transparent species of teleost fish closely related to zebrafish (*Danio rerio)*. Males produce bursts of pulsatile, click-like sounds and a distinct postural display, extension of a hypertrophied lower jaw, during aggressive but not courtship interactions. Females lack a hypertrophied lower jaw and show no evidence of sound production or jaw extension in such contexts. Novel pairs of size-matched or -mismatched males were combined in resident-intruder assays where sound production and jaw extension could be linked to individuals. Resident males produce significantly more sound pulses and extend their jaw more often than intruders in both dyad contexts, and relatively larger males are significantly more sonic and exhibit more jaw extensions in size-mismatched pairs. The majority of highest sound producers in both contexts also show increased jaw extension during periods of heightened sonic activity. These studies firmly establish *D. dracula* as a sound-producing species that modulates both acoustic and postural displays during aggressive interactions based on either residency or body size, providing a foundation for further investigating the role of multimodal displays in a new model clade for neurogenomic studies of aggression, courtship, and other social interactions.

## INTRODUCTION

Animals across taxa have distinct behavioral repertoires utilizing a variety of signaling channels and modalities to mediate complex social interactions (Bradbury and Vehrencamp, 2011). Signal structure within a modality can further vary at a species or individual level depending on the environment and social context (Bradbury and Vehrencamp, 2011). Acoustic communication provides many such examples (e.g., Gerhardt and Huber, 2002), with signals differing in parameters ranging from amplitude, duration and fundamental frequency to the time interval between repetitive components within a call and between calls (e.g., Davies and Halliday, 1978; Clutton-Brock and Albon, 1979; Sebastianutto et al., 2008; Odom et al., 2021). Acoustic repertoires also often differ between the sexes (Gerhardt and Huber, 2002; Bradbury and Vehrencamp, 2011). In many species of teleost fish, males often exhibit more diverse acoustic repertoires than females, although detailed investigations of female sound production are scarce (Amorim, 2006; McIver et al., 2014; Pereira et al., 2014; Amorim, 2015; Ladich and Maiditsch, 2017). Sex differences in soniferous behavior often reflect underlying structural dimorphisms that are frequently associated with hypertrophied muscles driving swim bladder vibration or stridulation of pectoral fin rays that lead to sound production (Rice et al., 2022). From an ecological perspective, sound production can figure prominently into aggressive competition for mates, nest sites, shelters, and territories (Andersson, 1994). Sonic signaling can enable contests to be resolved through less costly measures before escalating to more intense stages of engagement involving direct contact and possible physical damage (e.g., Davies and Halliday, 1978; Clutton-Brock and Albon, 1979, Hsu et al., 2008; Oliveira, Silva and Simo, 2011; Green and Patek, 2018). In so doing, these signals may communicate information about a contestant’s identity (e.g., age and sex), motivation to fight for a resource item, which is often associated with higher signaling display rate (Burmeister et al, 2002; Arnott and Elwood, 2007; Triefenbach and Zakon, 2008), and physical attributes such as body size, which can figure prominently into determining the outcome of aggressive conflicts (e.g., Davies and Halliday, 1978; Clutton-Brock and Albon, 1979; Bradbury and Vehrencamp, 2011; Amorim, 2015; Conti et al., 2015; Billings, 2018).

Our main intent here is to provide a behavioral baseline for studies of acoustic communication during social interactions in *Danionella dracula*, a miniature species of cypriniform fish (Britz and Conway, 2016) that we report are sonic and can be readily bred in captivity. We describe sound production in *D. dracula* mainly within the context of aggressive interactions because we found that males produce relatively simple, broadband sounds apparently solely during aggressive but not courtship encounters; females appear to be silent during all such interactions. Given the close phylogenetic relationship of *Danionella* species to zebrafish (*Danio rerio*), they are also amenable to genetic manipulation using the zebrafish molecular toolbox (Schulze et al., 2018). *Danionella dracula* and its sister species further offer multiple advantages for behavioral neuroscience given their especially small size as adults (see below) and transparency into adulthood (e.g., Fig. 1A-C), thus facilitating non-invasive, optical imaging of the brain using multiphoton microscopy (Schulze et al., 2018, Chow et al., 2020; Akbari et al., 2020, Akbari et al., 2021). Despite possessing these attractive features for genetic and neural studies of social behavior, these behaviors have not yet been described in a comprehensive manner for any *Danionella* species.

**Fig. 1:**
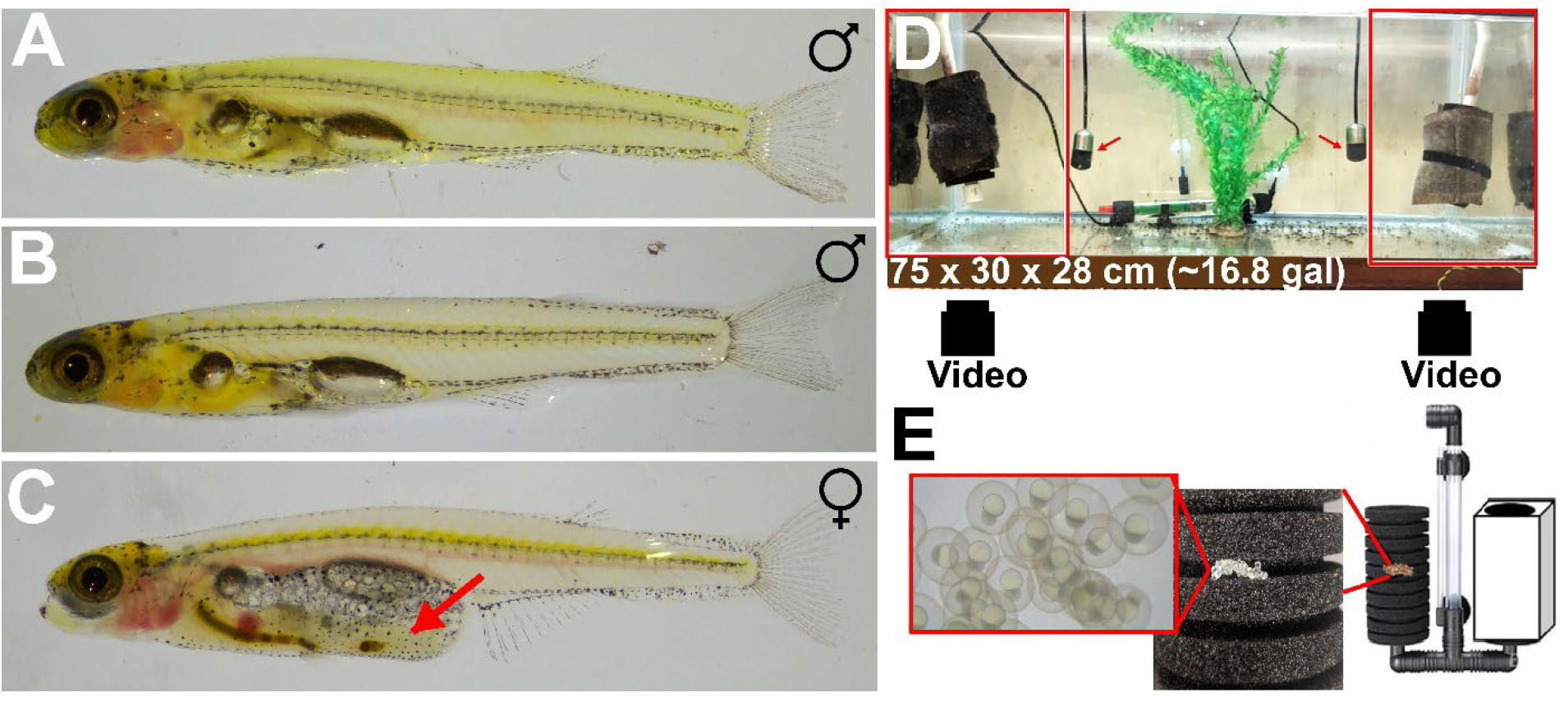
Male and female *Danionella dracula* and community tank. A. Adult male with green to yellow-green coloration, standard length (SL) = 17.2 mm B. Adult male without green coloration, SL = 16.8 mm. C. Adult female without green coloration, SL = 17.1 mm. Ovary indicated by red arrow. D. Community tank setup (details in Materials and Methods); red boxes at either end indicate video camera field of view. External video cameras indicated by black boxes below tank. Red arrows indicate hydrophones. E. Eggs in nest site: insets from left to right show developing eggs, egg cluster in crevice of sponge filter nest site, and schematic of nest site used for egg collection.

*Danionella dracula* was first described in 2009, known only from a small stream in the Kachin state in northern Myanmar (Britz, Conway and Rüber, 2009, Britz and Conway, 2016, Britz, Conway and Rüber, 2021). Nothing is known regarding the specifics of its behavior in the wild, largely because its natural habitat is currently relatively inaccessible due to political unrest (e.g., see Goldman, 2021; UN Human Rights Council, 2018). Like other members of the genus, *D. dracula* exhibits developmental truncation, as they retain some larvae-like traits as miniature adults (paedomorphy) and only reach 12-18 mm in standard length (Britz, Conway and Rüber, 2009; Britz and Conway, 2016; Conway, Kubicek, and Britz, 2021; this report). Morphological studies predicted a sex difference in soniferous behavior for *Danionella* species based on documentation of a putative sonic “drumming apparatus” in adult males, but not females (Britz, Conway and Rüber, 2009, Britz and Conway, 2016, Britz, Conway and Rüber, 2021, B. Judkewitz, personal communication). Sound production was reported earlier for male *D. cerebrum* (Schulze et al., 2018; initially designated *D. translucida*, see Britz, Conway and Rüber, 2021). Unlike other species within the genus so far described, *D. dracula* has a hypertrophied lower jaw and a series of odontoid processes that resemble fangs (Britz and Conway, 2016).

We report the establishment of a laboratory-based breeding colony of *D. dracula* to study courtship and aggressive behavior over the use of nest sites. Furthermore, we present the results of a resident-intruder assay to examine acoustic and postural signaling during dyadic aggressive interactions in size-matched and size-mismatched male encounters. Previous research in aggression has utilized behavioral tests like resident-intruder assays (see Koolhaas et al., 2013) and other dyadic designs to study how two individual males can compete over a resource and what signaling parameters correlate with differences in the ability for an animal to win a contest, i.e., the animal’s resource-holding potential (RHP; see Bradbury and Vehrencamp, 2011). To investigate the effect of two factors that contribute to fighting ability, residency status and relative body size (see Hack, Thompson and Fernandes, 2010; Jennions and Backwell, 1996; Jackson and Cooper, 1991), on agonistic signaling behaviors in *D. dracula*, we focused our quantitative analyses on two prominent display characters that we show distinguish *D. dracula* from others within the genus: temporal features of acoustic signals and extension of the hypertrophied lower jaw. Our original intent was to only investigate acoustic displays. However, we noticed early on that males often extended their lower jaw during sound production. This suggested that sonic activity may depend biomechanically on jaw extension, reminiscent of the involvement of jaw movement in clownfish (*Amphiprion clarkii*) sound production (Olivier *et al.*, 2015). Thus, we hypothesized that the amount or temporal pattern (e.g., inter-pulse interval) of sound production and jaw extension by an individual male would be influenced by residency and/or relative size.

## MATERIALS AND METHODS

### Colony formation and breeding

Adult *D. dracula* were originally purchased from commercial dealers (The Wet Spot Tropical Fish, Portland, OR; Invertebrates by Msjinkzd, York, PA) and bred in environmental control rooms at Cornell University, Ithaca, NY. Fish were kept at water temperatures of 23-25.5° C and on a 16:8 light: dark cycle. We found that *D. dracula* breeds best with direct overhead lighting. Fish were kept in rooms with artificial or mixed natural and artificial overhead lighting. There were no obvious differences in social behavior, breeding, or rates of development between the two lighting conditions. All dyad contests were conducted in rooms supplemented with artificial lighting on a 16:8 light: dark cycle. All animals were fed twice a day, *Artemia nauplii* in the morning and ground fish flake (TetraMin Tropical Flake) in the evening. Plastic plants were added to each tank for fish to acclimate to the aquarium setting. We also found that *D. dracula* is a diurnal species, with more sonic activity during daylight hours, like their sister species, *D. cerebrum* (Schulze et al., 2018). All behavioral observations for colony, community, and dyad tanks were carried out between 09:00 – 17:00 h. All procedures were approved by the Institutional Animal Care and Use Committee of Cornell University.

Fish were bred in 2.5-125 gal aquaria that housed populations of varying density based on tank size, in ratios of 1:2 males: females, with at least three males per tank (Fig. 1D). Nest sites were made from double sponge water filters (XY-2822 Air Pump Double Sponge Water Filter, Xinyou) that contained nine 4-mm crevices for spawning (Fig. 1E). The sponges were placed at opposite ends of the tank (Fig. 1D) and covered with a BIO-CHEM ZORB filtration cartridge (API Fishcare CRYSTAL Filtration Cartridge), as it is required for the crevices to be enclosed for breeding (Fig. 1E). Eggs were collected daily by removing the nest, unrolling the filtration cartridge and gently moving the clusters of eggs to acrylic cylinders (10 cm diameter) with a mesh bottom that rested within a 50-gal aquarium (see eggs in Fig. 1E). Larval fish were fed AP100 Dry Larval Diet (Zeigler Bros, Inc.) twice a day for 10 d in these smaller cylinders before being moved to 5-gal tanks where they were fed adult diet. Larvae became adults in 3 mos, visually determined by the presence of eggs in the abdomen of females and the hypertrophied jaw in males.

### Community tank observations

A 16.8-gal community tank with dimensions 75 x 30 x 28 cm was set up using the same parameters for colony system tanks to allow for behavioral observations in a reproductive context (Fig. 1D). A heater kept the tank at 25°C. Males (3) and females (6) were placed into the tank 30 mins after dye labeling. Each male had a muscle segment labeled in the tail with either red, green, or black dye to allow three independent observers to determine identity while watching the tank and in video recordings (Tissue Marking Dye Kit, MDT100, Sigma-Aldrich). Females were also labeled using the same method, thus all fish in the tank went through the same injection process. Sounds were recorded with a hydrophone (Aquarian Audio H1a) placed next to each nest site and connected to a 30-fps video camera (Canon Vixia HFR500) using a mono to stereo adaptor to synchronize the audio collection with the video.

Fish were allowed an acclimation period, which concluded after one week at the onset of courtship. Then, three independent observers used the software BORIS (Friard and Gamba, 2016) to conduct focal sampling and live observation, alongside collecting video and acoustic data centered on the two nest sites, to observe each of the three labeled males. Behavioral observations were based on three randomly selected, 15-min periods, made up of three 5-min periods where each of the different focal males were observed, between 09:00 and 17:00 for each of seven days. Fish received the same diet regime as colony tanks, with the first feeding at 10:00-10:15 and the second at 17:00. Live observations were synchronized to the video with a red LED pressed at the start of the observation period by the observer. Observers sat 46 cm in front of the tank and used keystrokes to signify behavioral events of interest in the focal male (Table S1). These characterizations were verified with the video camera and sound data collected.

### Resident-intruder assays

Males were removed from colony tanks 22 h before the resident-intruder assay. The resident male was housed in the experimental rectangular tank, which was 14 cm x 5 cm x 5 cm (Fig. 2A). The intruder male was housed separately in an 8 cm x 8 cm x 8 cm tank. Both tanks contained the same volume of water, 315 cm^3^, with a similar depth of 4.5-5 cm. Males were selected for the dyads to be as close as possible in size (standard length: 13.19-17.94 mm), with the relative size difference between males being less than 0.5 mm (0.2-2.9% relative size difference). Taking advantage of adult male color variation, which is not observed in adult females (Fig. 1A-C), residents and intruders were selected to be different colors in size-matched assays to make them readily obvious in the videos. One male was greener in coloration than the other, which was more pale yellow; half of the size-matched residents were pale yellow, and half were green (Fig. 1A, B). There was no apparent effect of color on total sound production in size-matched contests (F_1,85_=1.42, p=0.2361). However, color is among the many possible variables that could be controlled and/or manipulated in future studies, especially with a large sample size. Resident-intruder assays were also performed with size-mismatched dyads. Residency and relative size status were counterbalanced across dyads. In size-mismatched contests, males were easily distinguished from each other in the videos based on size, as one fish was distinctly larger than the other in each of the assays. The relative size difference between males in the dyads ranged from 1.6 to 4.4 mm (standard length larger fish: 14.64-17.81 mm; smaller fish: 12.08-13.86 mm; 11-28% relative size difference). Fish were fed their community tank diet in the evening and morning before testing. Both tanks were aerated with an air stone, and water was novel system water that had not housed adult fish previously. Three of the four tank walls were black and opaque, allowing for better contrast for later fish identification. The experimental tank used to run resident-intruder trials alone contained a hydrophone (Aquarian Audio H1a). The size of the hydrophone was chosen to best resemble a *D. dracula* nest site; this type of hydrophone has been observed to elicit crevice-seeking behaviors in males. Following a 22 h acclimation period lasting from 13:30 on day 1 until 11:30 on day 2, each resident-intruder trial lasted for 2 h, beginning at 11:30 (Fig. 2B). The intruder male was added to the experimental tank with a small net to begin the resident-intruder assay (Fig. 2B, C). Sound production and video were recorded through the hydrophone attached to a 30-fps video camera (Canon Vixia HFR500). Two of the 10 size-matched dyads were removed from analysis as the fish did not acclimate, swimming continuously against the sides of the miniature tank. Two of the 10 size-mismatched dyads were removed from analysis. In one dyad, both fish were swimming continuously against the sides of the miniature tank. In the second dyad, both fish were intensely engaged in an escalated aggressive interaction making it impossible to ascertain the identity of the sound producer (similar to extended aggressive interactions in community tanks). Therefore, 8 dyads were analyzed for each context, size-matched and mismatched.

**Fig. 2:**
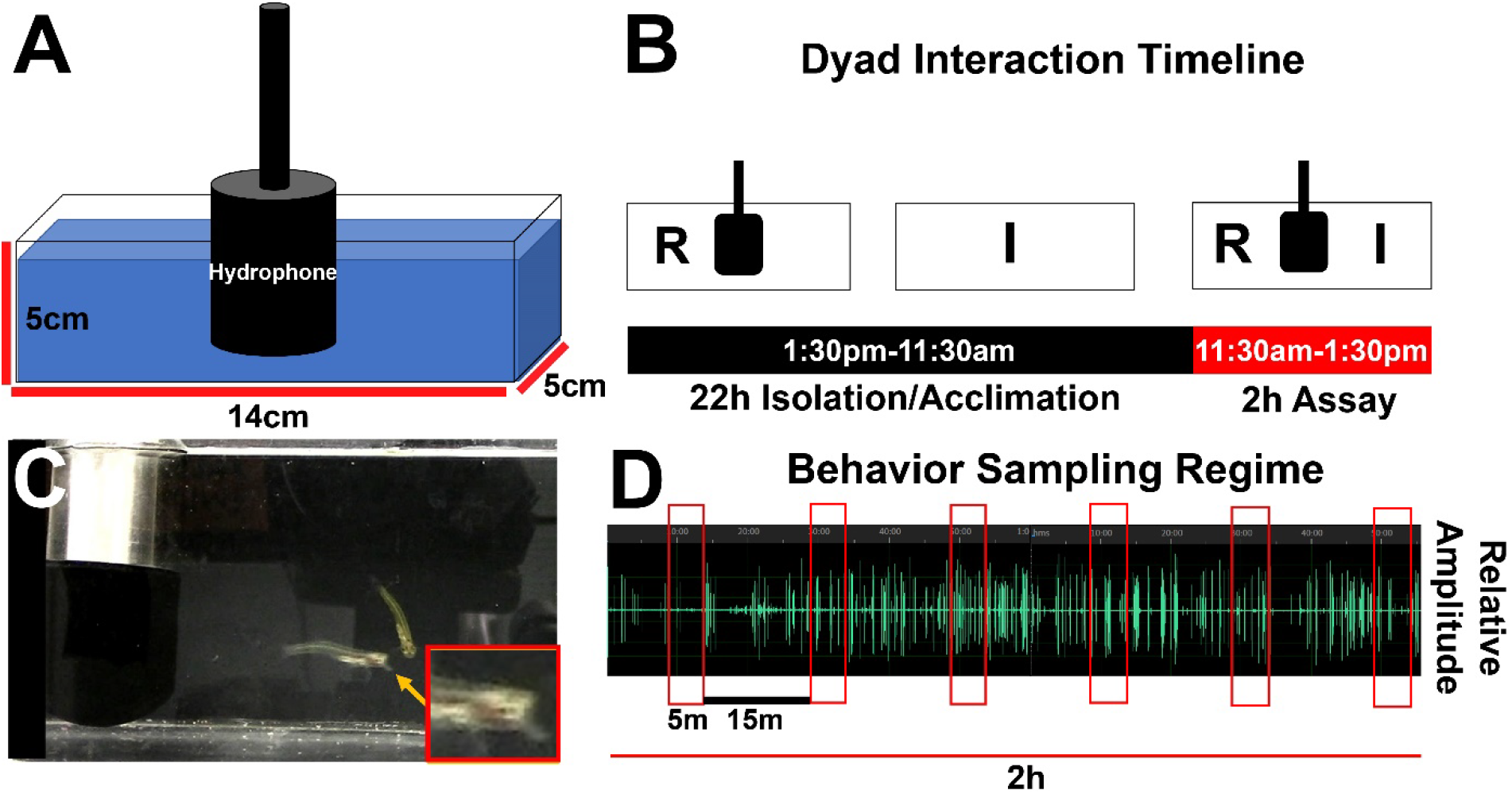
Dyad testing tank for male *Danionella dracula*. A. Experimental tank schematic; dimensions:14 cm x 5cm x 5cm. Black cylinder represents the hydrophone. B. Dyad interaction schematic and timeline. R stands for Resident and I for Intruder male. Black indicates the 22 h isolation/ acclimation period (1:30pm-11:30am) that fish undergo before the 2 h assay (11:30am-1:30pm), which is indicated in red. C. Two males in testing tank, one with jaw extended as it lunges at the other. Extended jaw indicated with orange arrow; insert is closeup of anterior body region and jaw. D. Behavior sampling regime. Red boxes on oscillogram (sounds in green) indicate six 5 min time periods sampled at 15 min intervals (black line) over 2 h assay.

Five single-male control trials were conducted following the same procedure, but only one male was placed in the experimental chamber and an intruder male was not added to the chamber at the start of the 2 h trial. Five additional control dyads were conducted using male-female pairs. Five males and five females (standard length: males, 14.06-17.08 mm; females, 13.73-17.12 mm) were combined in single pairs in the experimental tank following the same procedure, varying in size differences between males and females (0.3-3mm; 2-20% relative size difference). Two of the residents were male and three were female.

### Audio recordings and analysis

Hydrophone recordings of resident-intruder trials lasted for the full two hours. Due to the large number of sounds observed in the 2 h period, the oscillogram from each hydrophone recording after the initial acclimation period was split into six 5 min long time-periods separated by 15 min intervals (Fig. 2D). Sound characteristics in Table 1 were measured using Raven Pro 1.6 (K. Lisa Young Center for Conservation Bioacoustics, 2021) Recordings typically displayed a high signal to noise ratio (e.g., Fig. 2D). Pulse duration was determined by first measuring the maximum amplitude of an individual sound pulse’s waveform. This value was then divided by 4 to get a quarter amplitude value, and the pulse duration was determined as the duration of the pulse where waveform peaks were all greater than the quarter amplitude value. Pulse peak frequency was measured for each pulse after selecting the pulse using the pulse duration criteria above, using Raven’s Peak Frequency measurement. This measurement in Raven is calculated from the spectrogram of the sound and is the frequency at which the maximum/peak power occurs within the selection. On the recording, we observed individual pulses occurring very close to each other in time, forming apparent clusters composed of multiple pulses. We measured the duration of time between all neighboring individual pulses, the inter-pulse interval (IPI), and pooled all the male IPI data together. We used the mode value of 34 ms (Fig. S1) to set boundaries for individual burst types, where pulses that composed a burst had to possess an IPI within two standard deviations of the mode, or be less than 70 ms apart. This criterion allowed us to identify burst types ranging up to 6 pulses in length. All IPIs greater than 70 ms were defined as inter-burst intervals (IBI), i.e., the duration of time between bursts composed of multiple pulses.

**Table 1.** Sound Characteristics and Definitions in *D. dracula*

| Sound Characteristic | Definition |
| --- | --- |
| Pulse Duration | Total duration of one individual sound pulse. |
| Peak Pulse Frequency | Frequency at which the maximum peak power occurs within one individual sound pulse. |
| Burst | A train of individual pulses (1-6 observed), where pulses are separated by $\leq 70$ ms. |
| Inter-Pulse Interval | Time between individual pulses in a single burst of pulses, where pulses are separated by $\leq 70$ ms. |
| Inter-Burst Interval | Time between individual bursts, where interval between two successive pulses is $\geq 70$ ms. |

To characterize the amplitude range of *D. dracula* sound pulses, we recorded sound production using a calibrated hydrophone (8013, Brüel & Kjaer) connected to a conditioning amplifier (2635, Brüel & Kjaer) captured on a digital recorder (LS-12, Olympus). We first recorded sounds in a large community tank (122 x 46 x 74 cm, ~100 gal) with 75 fish and 4 nest sites (one in each corner of the tank). The hydrophone was suspended 15 cm beneath the water surface and equidistant (15.24 cm) between two nest sites on the left side of the tank. We analyzed sounds from 2 h of audio recorded 10:00-12:00. We next recorded sounds in the resident-intruder assay in an acoustic isolation chamber (Industrial Acoustics). The hydrophone was positioned 2 cm from the right side of the rectangular experimental tank. Two size-matched males were introduced to the tank as previously described and 1 h of audio was captured. Sound pulses were isolated and analyzed in Raven Pro 1.6 using a custom script.

### Video analysis of sound production and jaw extension behavior

For the resident-intruder assays, an observer blind to resident-intruder status verified instances of male sound production by watching the synchronized video at 0.3X the normal speed. Sound bursts were attributed to an individual male based on associated lunging movement. This association between lunging and sound production was established based on our analysis of a 4 min portion of a 23 min recording of a *D. dracula* male continuously lunging at its reflection in the wall of the tank (Fig. 4B, oscillogram in C, Movie 1) For this recording, an observer first coded in BORIS all lunges directed at the reflection, without sound. The time point of an instance of lunging was determined as the first frame where the male fish oriented its head towards and swam rapidly towards its reflection. Burst and pulse start times from the same portion of the recording were measured using Raven Pro 1.6, so the time between a lunge and a burst could be examined for temporal proximity, as is described in the results (see Fig. S2A).

Observers distinguished fish in the size-matched assays based on coloration: green or pale yellow (see above), as well as other identifying features such as body girth. In size-mismatched assays, one fish was distinctly larger than the other and the two fish could be readily identified based on relative size. Coding of jaw extension was done by a third observer in BORIS who watched the video at 0.3X the normal speed without sound. The time point of an instance of jaw extension was determined as the first frame where the lower jaw was first extended from the head.

### Statistical analyses

Statistical analyses were performed in R version 4.1.1 using the lme4 package (R Core Team, 2021). We investigated whether agonistic behaviors varied between residents and intruders across each behavioral context. To do this, we ran a repeated measures Linear Mixed Model (LMM) with pair identity as a random effect to examine if agonistic behaviors differed between residents or intruders in the size-matched context, and whether this usage may change across the length of the behavioral trial (e.g., time-periods 1-6 of behavioral sampling; see above for more details about sampling). As in the previous models, we also investigated if overall behavior usage varied between residents and intruders, and large and small fish in size-mismatched contests, including both residency and size as predictors and an interaction term for residency and size.

To test the hypothesis that multi-pulse burst usage may vary between residents or intruders in the different contexts, we reduced each animal’s multi-pulse bursts (2 – 6 pulses/burst) into a single metric. As such, we performed a principal component analysis (PCA), using the *psych* package and *principal* function. In these models, we entered the number of different burst types for each male into R, and generated PCs separately for size matched and mismatched contexts. Higher PC scores represented greater multi-pulse burst usage. To next investigate whether PC1 (multi-pulse burst usage) varied between residents and intruders in the size-matched context, we used a LMM with pair identity as a random effect. In size mismatched contests, we also used a LMM to investigate whether multi-pulse burst usage differed between residents and intruders as well as large and small individuals’ size, by including both residency and size as predictors and an interaction term for residency and size. Finally, using total sound production as a continuous variable, we tested whether overall sound production was related to multi-pulse burst usage (PC1) by using a linear model separately for size-matched and - mismatched contexts.

## RESULTS

As stated in the Introduction, our main goal here is to provide a behavioral baseline for studies of acoustic communication during social interactions in *Danionella dracula*. For general context, we first provide qualitative descriptions of courtship, spawning and male aggressive behavior in a breeding colony of *D. dracula*, the first known laboratory colony of this species. We focus on males, as we found that they are the apparent sonic sex in this species and apparently only make sounds during aggressive encounters. Then, we provide quantitative assessments of acoustic signaling during resident-intruder male interactions, including a description of spectral and temporal properties of sound pulses. We also assess a prominent postural display, extension of the hypertrophied lower jaw, which is an additional display behavior that we found occurs along with sound production during aggressive interactions and distinguishes *D. dracula* from other species within the genus.

### Courtship and spawning

*Danionella dracula* readily bred and produced courtship and aggressive behaviors in laboratory colony and community tank settings. Although we focus on male aggression in this report, we first provide a representative description of courtship and spawning in a colony tank, where we first observed these behaviors, for broader context here and in future studies. No sound production was ever recorded during either courtship or spawning in our colony and community tanks. Males typically swam from nest sites to court females who were often schooling around the tank. Fig. 3A-F show single frames from a video record of courtship and spawning in the presence of a black vertical tube that was originally part of an underwater filtration system in a colony tank (also see Movie 2). The male approached a female and swam beneath her, directly hovering beneath her egg vent (Fig. 3A, B). He then rapidly moved his pectoral fins and vibrated his body and head back and forth beneath the female. Next, the male swam back to the nest site that he swam closely around during the day. The female swam behind the male to the same nest (Fig. 3C) and the male then entered the spawning crevice head-first (Fig. 3D) followed by the female (Fig. 3E, F). After spawning, the female left the nest and the male emerged from the nest. He then swam closely around the nest, which we refer to as nest circling, and courted additional females throughout the day.

**Fig. 3:**
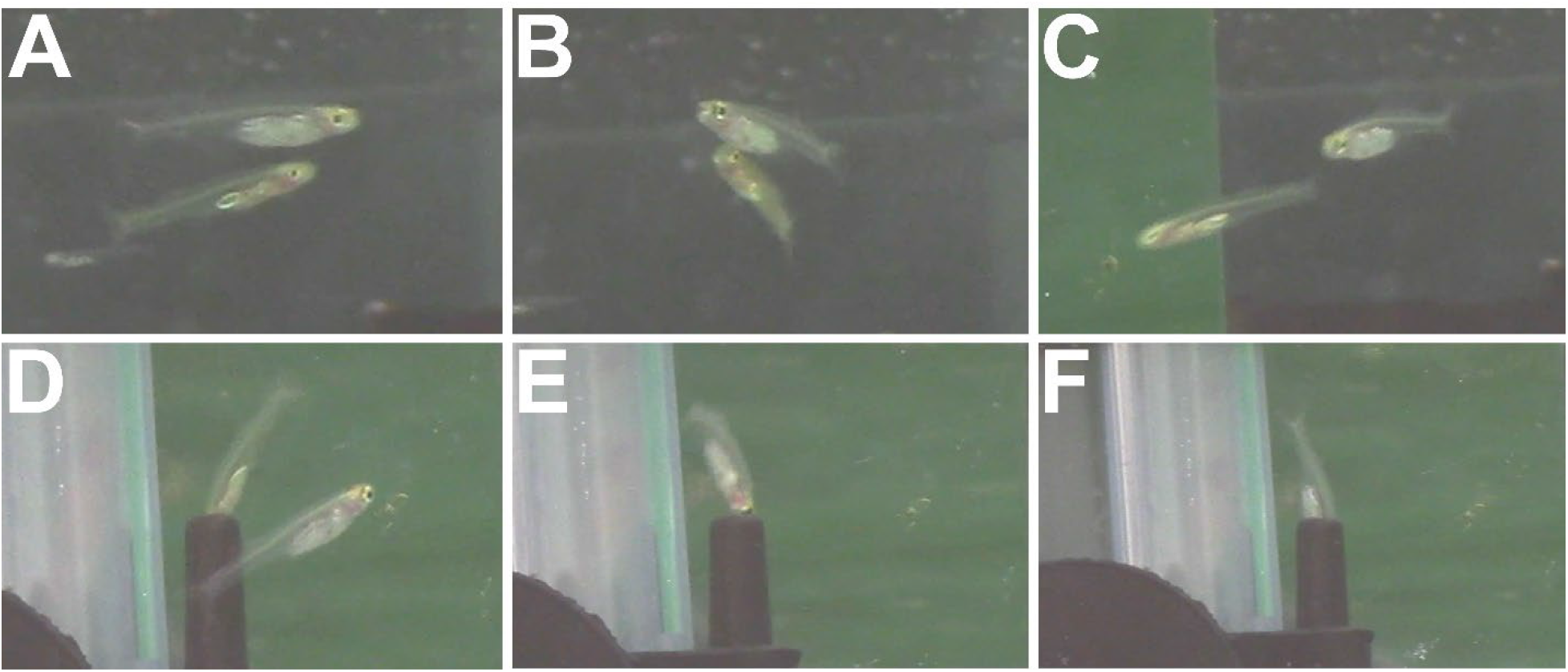
Video frames of *Danionella dracula* courtship behavior in colony tank. A. Male swims below female B. Male vibrates his body back and forth beneath female egg vent. C. Female follows male back to nest entry crevice (part of an underwater filter). D. Male enters headfirst into crevice E. Female orients head towards crevice. F. Female swims headfirst into crevice.

Spawning cannot be directly observed in either colony or community tank settings as it only occurs in enclosed crevices. It is inferred to take place in the crevices for two reasons. First, clutches of 12-24 eggs are found in the crevices of the artificial sponge nests; eggs are found in nest grooves at any time of the light cycle following spawning interactions (see Fig. 1E). Second, we often find a male and/or a female on top of a cluster of eggs within the grooves of a sponge nest during daily nest checks.

### Male aggressive displays

In colony tanks, we often observed males swimming throughout the day around artificial nest sites that are dark-colored objects containing crevices for spawning (see Fig. 1D, E). By observing dye-marked males in a controlled community tank setting over a two-week period, we often recorded the same male swimming within 2-3 body lengths around the same nest site for 1-8 days, suggesting nest fidelity. These males also lunged at other males who swam within 2-3 body lengths around the nest site, often producing a burst of stereotyped sound pulses close to the same time as the lunge (Fig. 4A).

**Fig. 4:**
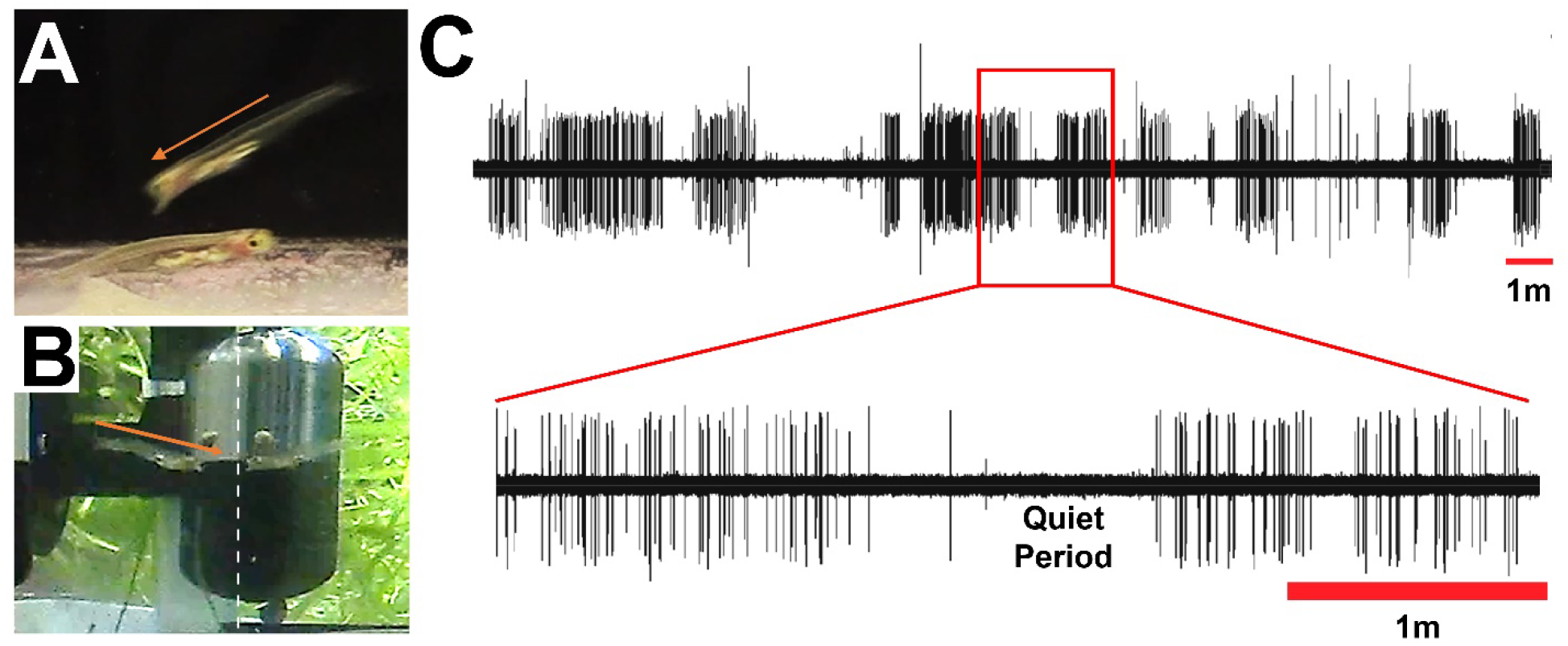
Video frames and oscillogram of *Danionella dracula* male-male aggressive interaction. A. Frame from resident-intruder assay of male lunging at other male and extending its lower jaw; direction indicated by orange arrow. Both resident and intruder males lunge and produce sound pulses. B. Frame from male lunging at its own tank reflection. Direction of lunge towards tank glass indicated by orange arrow extending above entire dorsum of lunging male. White dashed line indicates reflective surface of tank wall. C. Oscillogram from 23 min recording of male in ‘B’ lunging at its reflection in tank wall on two timescales. Lower trace is 4 min section of the recording to show individual sound pulses. “Quiet Period” indicates period when focal fish was not oriented towards its reflection and was swimming around nest.

We concluded that sound production is primarily coupled to lunges at intruder fish, or the visual presence of another male, in the contexts critically assessed in our study. We reached this conclusion after combining these colony tank observations with another colony tank video recording of a male lunging continuously against its own reflection at the same time as apparent sound production (Fig. 4B; oscillogram from recording in 4C, see Movie 1). For a representative 4-min portion of this recording where the male lunged continuously at his reflection, we found that sounds were produced when the male was within one body length of its reflection in the tank wall, with its head and body oriented and moving towards its reflection in the tank wall (Movie 1). By contrast, there was essentially no evidence of sound production in a portion of the recording when the male was swimming around the nest and not lunging at its reflection (labeled in Fig. 4C); the two pulses one can see during the quiet period are timed with the fish lunging at another fish (Fig. 4C). We pooled together all the times between lunges and sounds and found a mean value of 579±27 ms; 88.4% of sound bursts and lunges (n=107/121) were less than 600 ms apart, closely temporally related (Fig. S2A).

In colony tanks, we also observed extended aggressive interactions lasting up to 30 mins that escalated to include mutual lunging, extension of the lower jaw, and vertical lateral displays until one male fled. It was not possible to readily assign sound production to either male during such interactions.

### Sound production during resident-intruder assay

To characterize sound production during aggressive interactions, we examined dyads in a resident-intruder context (Fig. 2A). Single male controls did not produce sounds and swam around the experimental chamber. There was also no evidence in male-female dyads that females made any sounds similar to males, consistent with absence of the putative sonic “drumming” muscle in females (Britz and Conway, 2016). A hydrophone placed in a 20-gal tank of 10 female fish for 10 h also did not provide any evidence of sound production by females.

Dyad interactions appeared to be primarily one-sided, where one fish emerged as the more apparent aggressor towards the other; 94.7% of total sounds (2705/2858) were associated with one male’s distinct lunge at the other fish (see example in Movie 3). 5.3% of sounds were too ambiguous to associate a sound with one male over the other, and they were not included in our consequent analyses. Out of these ambiguous cases, sound pulses could not be attributed identity because both fish were oriented towards each other and lunging at the same time as sound production (3.0%, 87/2858); both were swimming next to each other closely with their heads oriented towards each other but with no distinct lunges during sound production (1.4%, 41/2858), or both were obscured by the hydrophone or by both being in a corner of the tank (0.9%, 25/2858).

Males produced trains of sound pulses composed of repetitive bursts of up to 6 pulses per burst (see Table 1 for definition). In the dyad assay tank, individual sound pulses had a mean peak frequency of 1988.6±14.0Hz and 1961.6±11.5 Hz in size-matched and -mismatched contexts, respectively (see Fig. 5A - example spectrogram in 5A). Mean pulse duration was 2.0 ± 0.02 ms and 2.6±0.03 ms for size-matched and -mismatched males, respectively (Fig. 5C). The mean IPI was 42.9±0.6 ms and 39.5±0.5 ms in size-matched and mismatched contexts, respectively (e.g., see 2- and 3-pulse bursts in Fig. 5B). In both dyad contexts, males primarily used single pulses (size-matched, 65.3% of all bursts; size-mismatched, 64.1%) and 2-pulse bursts (size-matched, 23.4%; size-mismatched, 27.1%) that overlapped with lunging at other males (Fig. 5D). Three and higher pulse bursts occurred more rarely, with 3-pulse bursts making up 8.1% and 7.9% for size-matched and -mismatched male dyads, respectively (Fig. 5D). Four pulse and higher burst types made up only 3.2% and 0.9% for size-matched and -mismatched males, respectively (Fig. 5D). Individual fish produced multiple types of bursts containing differing numbers of pulses throughout the interaction (e.g., see Fig. 5B). Mean IBI was 12.6±1.6 s and 10.5±0.7 s for size-matched and -mismatched contests, respectively.

**Fig. 5:**
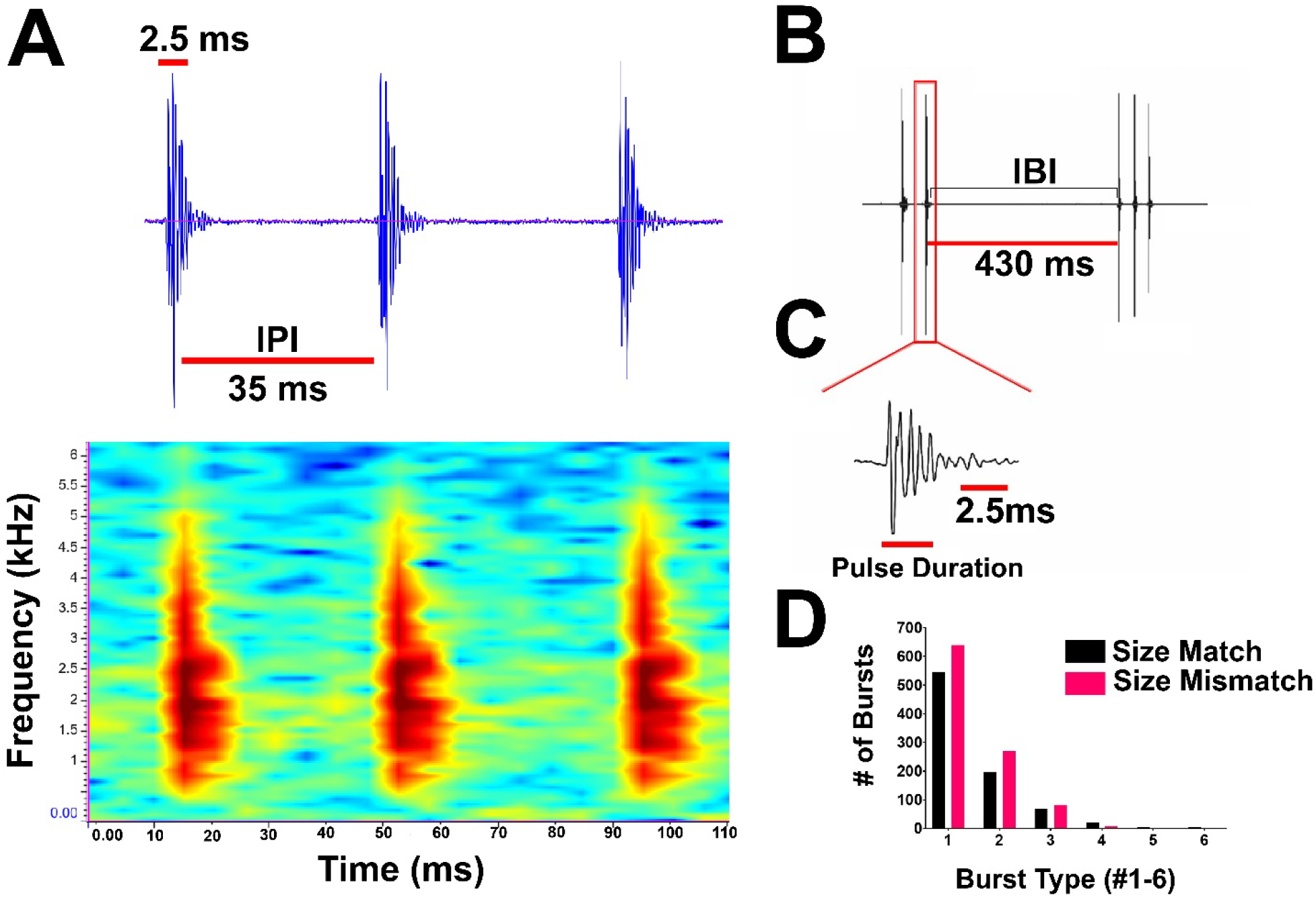
Multi-pulse sound bursts of male *Danionella dracula*. A. Top: Oscillogram of 3-pulse burst made by lunging male. Red lines indicate for this burst that pulse durations were close to 2.5 ms (top, see criteria in Materials and Methods) and pulses were separated by an inter-pulse interval (IPI) around 35 ms (bottom). Below: spectrogram of the same 3-pulse burst. Darkest red indicates peak power frequency, which in this burst was 1968.750 Hz. B. Example 2- and 3-pulse bursts separated by inter-burst interval (IBI) of 430 ms. C. One of the single pulses on expanded timescale from the 2-pulse burst in B. This pulse was 2.5 ms in duration. D. Bar graph showing number of different types of multi-pulse bursts observed in size-matched (black) and size-mismatched (salmon pink) dyads.

We did not measure sound amplitude in the assays given our inability to measure sounds from individual sound producing fish that were always moving around one hydrophone during these dyadic interactions in small volumes. In a separate set of tests designed explicitly to characterize the amplitude range of *D. dracula* sound pulses, we recorded 365 sound pulses in a 100-gal community tank, where we found an amplitude range of 141.5-153.09 dB re 1 μPa.

When recording males in the miniature resident-intruder assay tank, we found an amplitude range of 119.09-153.46 dB re 1 μPa from 451 pulses. The background noise (RMS) in the community tank was 137.81±0.46 dB re 1 μPa, mostly composed of low frequency noise (peak ~20Hz, all power below ~200 Hz; when filtered, background is 113.1±0.57 dB re 1 μPa). In the assay tank, the background noise (RMS) was 105.43±1.22 dB re 1 μPa.

Regarding total acoustic display rate, males varied widely in the total number of sound pulses directed towards another male, with some time periods for individual males having more sound production than others over the course of the 2 h dyad trial (see Fig. 6A-D). In size-matched assays, males who produced sounds above the median of observed male total sound production were categorized as high sound producing males, and those exhibiting below the median were categorized as low sound producing males (Fig 6 A-E). High sound producers ranged up to 318 total sound pulses in the 30 mins sampled over the 2 h dyad trial (Fig. 6E). In size-mismatched assays, we used the same criteria as with size-matched males and found that high sound producers ranged up to 428 total sound pulses (Fig. 6E). This variation was not explained by body size, as we did not see any apparent correlation between total sound production or any of the different characteristics of sound production described here with standard length (mm) or size difference (mm) between competitors in a dyad (Fig. S3).

**Fig. 6:**
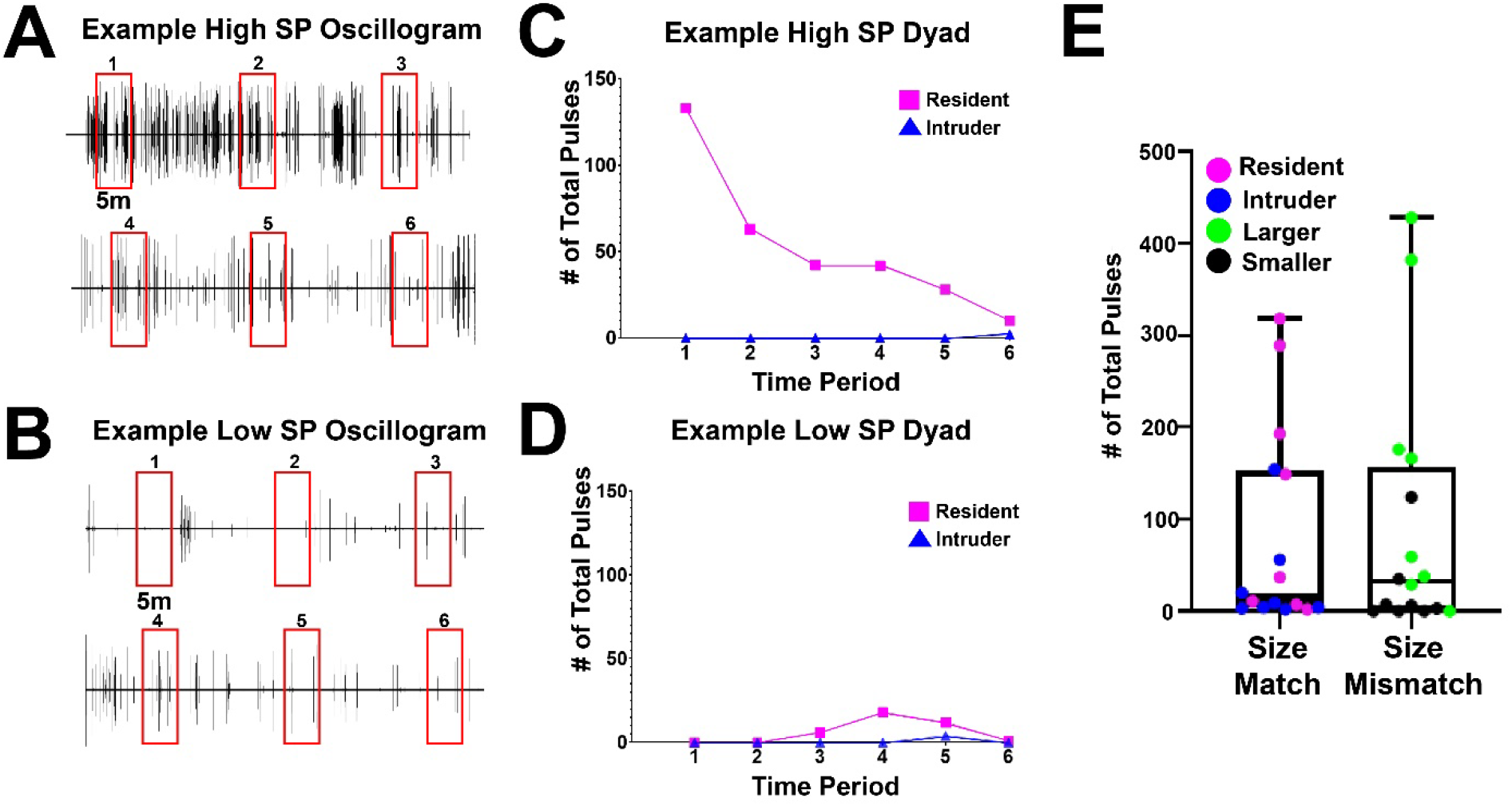
Total sound production (SP) by *Danionella dracula* males in dyadic contests. A. Representative oscillogram of high SP; rows 1 and 2 are from one continuous 2 h record. Red boxes indicate the six 5-minute sampled time periods (1-6) spaced 15 mins apart. B. Representative oscillogram of low SP; rows 1 and 2 are from one continuous 2 h record. Red boxes indicate the six sampled time periods. C. Example high SP from size-matched dyad in A, showing the total number of pulses for individual fish over the sampled time periods. Magenta squares indicate the resident fish’s SP values and blue triangles indicate the intruder fish’s SP values, which are at or near zero during all time periods. D. Example low SP from size-matched dyad in B, displayed as in C. Magenta squares indicate the resident fish’s SP values, which peaks at 20 sound pulses in time period 4, and blue triangles indicate the intruder fish’s SP, which in this case is also at or near zero during all time periods. E. Boxplots of total individual SP in size-matched dyads (left boxplot; magenta = resident fish, blue = intruder fish) and size-mismatched dyads (right boxplot; green = relatively larger fish, black = smaller fish). Black horizontal line within box indicates median that was used to designate fish as high or low sound producer.

### Jaw extension during resident-intruder assay

As noted in the Introduction, jaw extension often appeared to overlap sound production when we first observed *D. dracula* males in colony tanks, suggesting that sonic activity may depend on jaw extension. Thus, we also assessed the total instances of jaw extension in the dyads. Using linear regressions, we found that the number of sound bursts was significantly correlated with the number of jaw extensions in both size-matched and -mismatched contexts (F_1,14_=138.3, R^2^=0.9081, p<0.0001; F_1,14_=79.36, R^2^=0.8501, p<0.0001, respectively) (Fig. 7A, B). Jaw extension often, but not always (Fig. 7E), increased during the same 5 min time period samples of high sound production in both contexts (e.g., Fig. 7C, D).

**Fig. 7:**
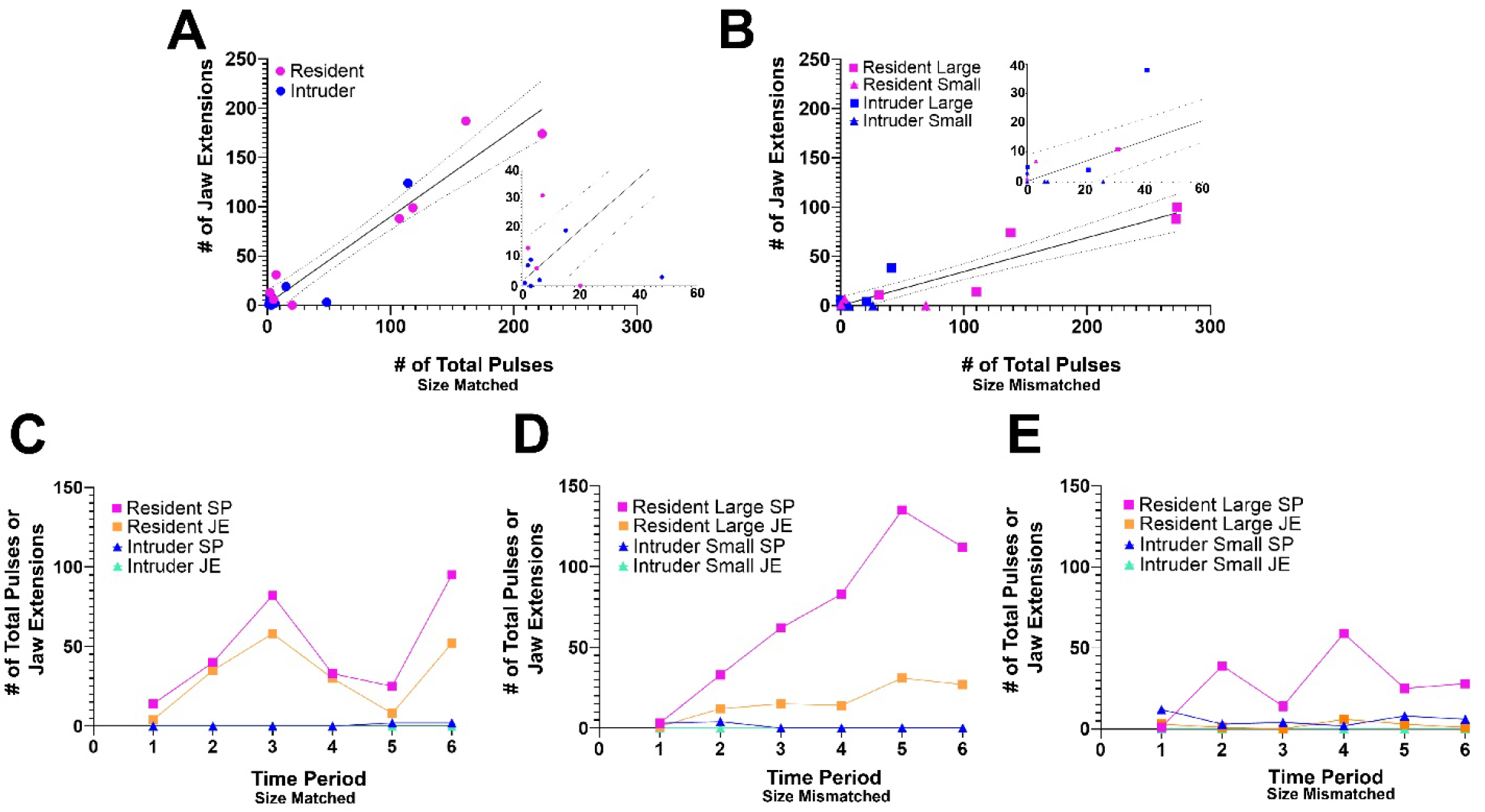
Temporal relationship between total number of sound pulses (SP) and jaw extensions (JE) for *Danionella dracula* males. A. Correlation plot of number of total sound bursts by number of total JEs in size-matched contests. Magenta circles indicate resident fish and blue circles indicate intruder fish. The black line indicates the linear regression, and the dotted light black lines indicate 95% confidence bands. Insert is closeup on origin of plot. B. Correlation plot of number of total sound bursts by number of total JEs in size-mismatched contests. Magenta squares indicate relatively larger resident fish and blue squares indicate relatively larger intruder fish. Magenta triangles indicate relatively smaller resident fish and blue triangles indicate relatively smaller intruder fish. The black line indicates the linear regression, and the dotted light black lines indicate 95% confidence bands. Insert is closeup on origin of plot. C-E. Examples of SP and JE for individual fish over the six sampled 5-min time periods from size-matched and - mismatched dyads. C. High SP and high JE from size-matched dyad. Magenta squares indicate the resident fish’s SP values and blue triangles indicate the intruder fish’s SP values. Orange squares indicate the resident fish’s JE values and light blue triangles indicate the intruder fish’s JE values. The intruder fish did not extend its jaw during time periods 1-6. D. High SP and high JE from size-mismatched dyad. Magenta squares indicate the relatively larger resident fish’s SP values and blue triangles indicate the relatively smaller intruder fish’s SP values. Orange squares indicate the larger resident fish’s JE values and light blue triangles indicate the smaller intruder fish’s JE values. The smaller intruder fish did not extend its jaw during time periods 1-6. E. High SP but low JE from size-mismatched dyad. Color code same as in D. The smaller intruder fish did not extend its jaw during time periods 1-6.

When directly examining the timing of the two behaviors to each other, we observed many instances where jaw extension appeared immediately before or after instances of sound production, as well as many instances of sound production without the presence of jaw extension, and vice versa. In aggregate, the evidence suggested that sound production and jaw extension were not obligatorily linked. Nonetheless, we wanted to more rigorously examine the temporal relationship between sound production and the start of jaw extension. We did this, in part, to provide baseline data for future studies assessing the impact of genetic manipulations on this temporal relationship, as well as for comparative studies of multimodal signaling between *Danionella* species. To maximize the sample size, these values were measured in males that exhibited both high sound production and jaw extension in size-matched (n=5) and -mismatched (n=3) contexts. For each dyad context, we pooled together all times less than 2 sec (considering events more than 2s apart to be independent) between sonic bursts and jaw extensions and found that mean time intervals between the two actions were 492 ± 20 ms and 590 ± 33 ms in the size-matched and -mismatched contexts, respectively (Fig. S2B, C). We found that the temporal separation between sounds and jaw extensions directed by the high sound producing and jaw extending male toward the other male could occur over a wide range, although a large number occurred within less than 100 ms in both dyad contexts and within as little as 2.5 ms and 1.3 ms of each other in size-matched and size-mismatched dyads, respectively (Fig. S2B, C). These results provided convincing quantitative evidence that sound production and jaw extension were independent events, although temporal gaps between the two actions could be very brief.

### Effects of Residency Status and Size on Display Rate

We next investigated the effects of residency status and size on sound production and jaw extension by examining the effect of being the resident or intruder in both dyad contexts, and the larger or smaller fish in size-mismatched assays. Residents produced significantly more sound pulses than intruders in size-matched (F_1,77_=10.19, p=0.002) (Fig. 8A) and size-mismatched (F_1,76_= 19.36, p<0.001) (Fig. 8B) contexts; larger males, irrespective of resident or intruder status, produced significantly more sound pulses than smaller ones (F_1,76_=15.38, p<0.001) (Fig. 8C). The interaction between residency and size was not significant (F_1,6_= 3.10, p=0.13). There was also no significant effect of time period on the total amount of sound production in size-matched (F_5,77_=0.46, p=0.81) and size-mismatched (F_5,76_=1.63, p=0.16) contexts, indicating there was not one specific time period (1-6) during the 2 h trial that fish across the dyads used increased sound production or jaw extension. The resident fish also extended its jaw more frequently than intruders in size-matched and -mismatched contexts (F_1,85_=5.39, p=0.02 and F_1,76_=9.10, p=0.003 respectively) (Fig. 8D, E). The same pattern held for larger and smaller fish (F_1,76_=23.42, p<0.001) (Fig. 8F). The interaction between residency and size in size-mismatched contexts was not significant (F_1,6_=2.25, p=0.18).

**Fig. 8:**
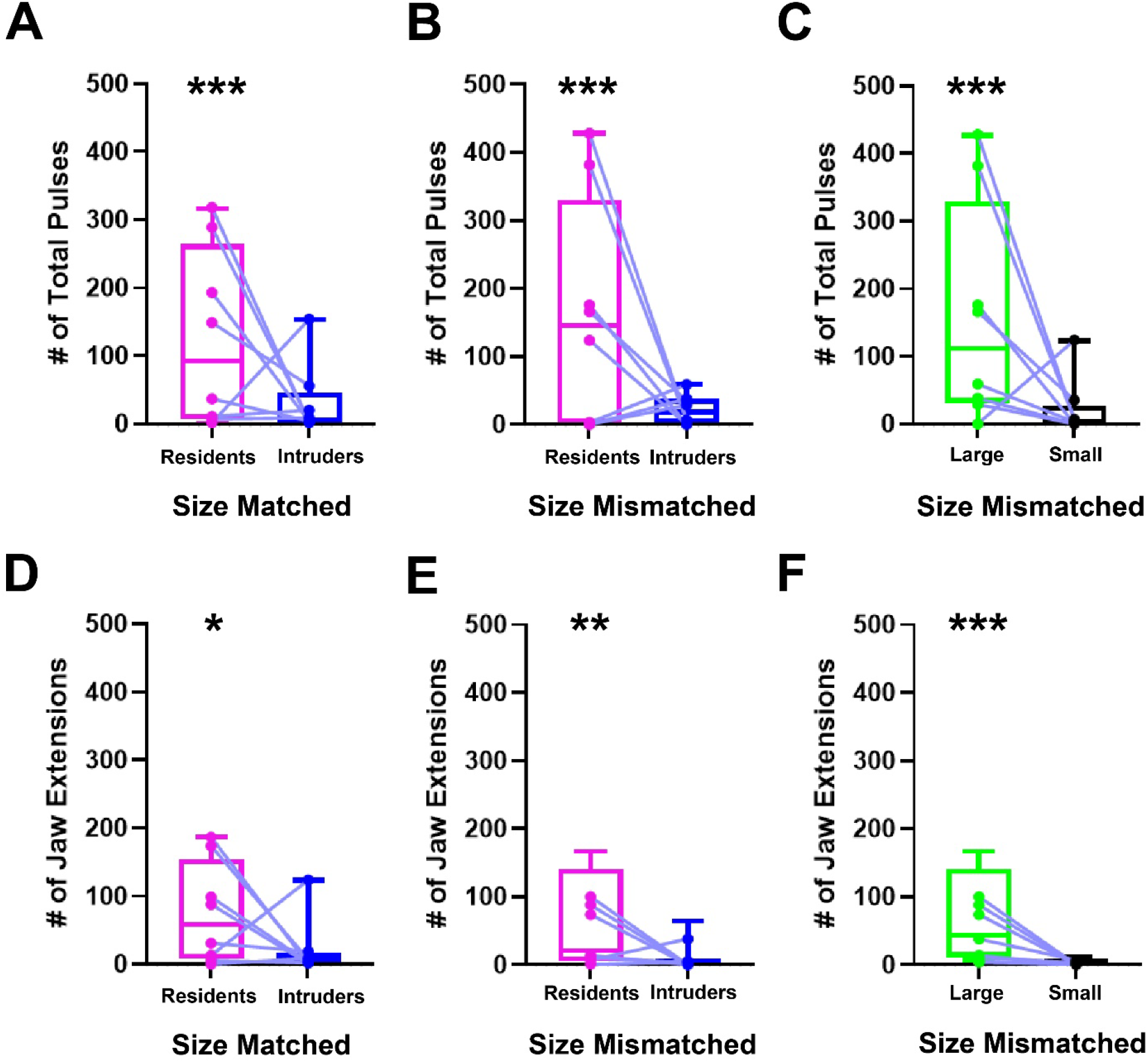
Fish identity and sound production for *Danionella dracula* males in dyadic contests. Boxplots comparing total amount of sound production (SP) as total pulse number (A-C) and total jaw extension (JE) behaviors (D-F) in size-matched and size-mismatched contexts. Individuals in the same dyad are connected between the two boxplots compared with a light blue line. * indicates significance as p compared to 0.05, ** p compared to 0.01, and *** p compared to 0.001. A. Boxplots of resident (left, magenta) fish SP and intruder SP (right, blue) in size-matched contests. B. Boxplots of resident (left, magenta) and intruder (right, blue) fish SP in size-mismatched contests. C. Boxplots of larger (left, green) and smaller (right, black) fish SP in size-mismatched contests. D. Boxplots of resident (left, magenta) and intruder (right, blue) fish JE in size-matched contests. E. Boxplots of resident (left, magenta) and intruder (right, blue) fish JE in size-mismatched contests. F. Boxplots of larger (left, green) and smaller (right, black) fish JE in size-mismatched contests.

### Effects of Residency Status and Size on Multi-Pulse Burst Usage

Beyond investigating the effects of residency and relative size on total display rate, we tested the hypothesis that there were differences in the temporal properties of the acoustic displays themselves, specifically in multi-pulse burst usage. We used a principal components analysis where the number of different burst types per male were entered as different variables (2-6 and 2-5 pulse bursts for each male in size-matched contests and size-mismatched contests, respectively; mismatched fish did not produce 6-pulse bursts). A single principal component (PC1) explained 77% of the variation of male usage of multi-pulse bursts in size-matched males and 65% of the variation in size-mismatched males (Table S2; Fig. 9A-E). In both models, higher PC1 scores represent animals that produced a greater number of multi-pulse bursts, often with these animals also using different types of bursts (2-6 pulse bursts).

**Fig. 9:**
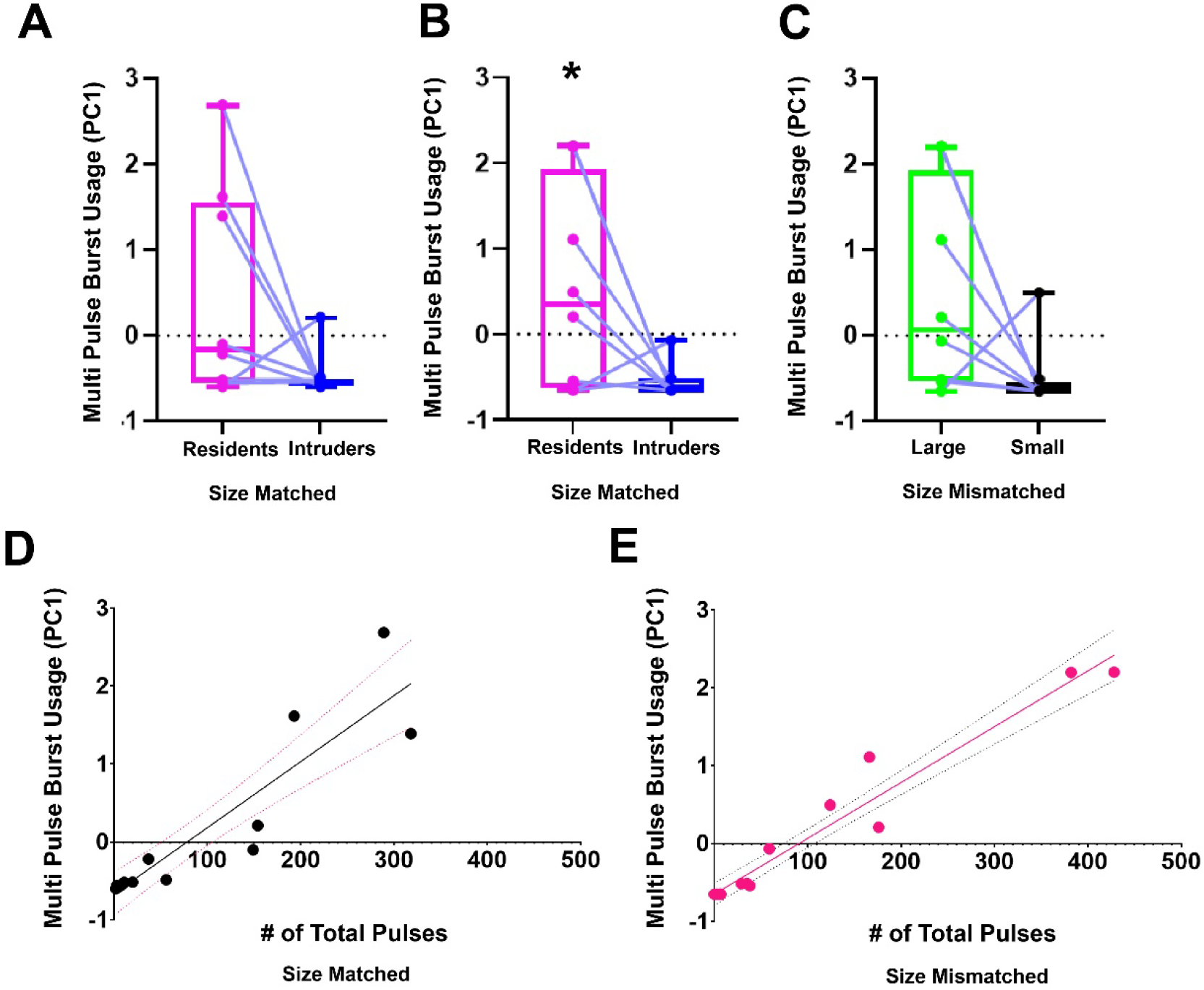
Multi-pulse burst usage for *Danionella dracula* males in dyadic contests.A. Boxplots of principal component 1 (PC1) describing individual male multi-pulse burst usage in size-matched contests. Residents are in magenta, left and intruders in blue, right. Individuals in the same dyad are connected by light blue lines. B. Boxplots of PC1 describing individual male multi-pulse burst usage in size-mismatched contests. Residents are in magenta, left and intruders in blue, right. Individuals in the same dyad are connected by light blue lines. * indicates significance as p compared to 0.05. C. Boxplots of PC1 describing individual male multi-pulse burst usage in size-mismatched contests. Larger fish are in green, left, and smaller fish in black, right. Individuals in the same dyad are connected by light blue lines. D. Correlation plot of number of total sound pulses by PC1 describing individual male multi-pulse burst usage in size-matched contests. Black circles indicate individual fish in assays. The black line indicates the fitted linear model and the dotted light pink lines indicate 95% confidence bands. E. Correlation plot of number of total sound pulses by PC1 describing individual male multi-pulse burst usage in size-mismatched contests. Salmon pink circles indicate individual fish in assays. The salmon pink line indicates the fitted linear model and the dotted black lines indicate 95% confidence bands.

We first investigated how multi-pulse burst usage (PC1) varied between resident and intruders across each behavioral context. In size-matched assays, residents and intruders did not significantly differ in the number of multi-pulse bursts (F_1,14_=4.09, p=0.06; Fig. 9A). Yet, in size-mismatched contests, residents produced significantly more multi-pulse bursts compared to intruders (F_1,12_=5.24, p=0.04; Fig. 9B). Larger fish in size-mismatched contests, regardless of residency status, did not significantly produce more multi-pulse bursts (F_1,12_=3.72, p=0.078; Fig. 9C).

The statistics of these comparisons did not indicate a strong predictive ability of residency or relative size status on multi-pulse burst usage in our assays. What we found to be more strongly predictive, however, is that when an animal had an increased acoustic display rate, as indicated by total number of pulses produced, they demonstrated higher multi-pulse burst usage. In a separate analysis, we investigated whether high sound producers produced more multi-pulse bursts compared to low sound producers by using total sound production as a continuous variable and running a linear model for each context. High sound production, regardless of dyad context, was significantly correlated with greater multi-pulse burst usage (size-matched: F_1,14_=71.71, R^2^=0.825, p<0.001; size-mismatched: F_1,14_=290.3, R^2^=0.9507, p<0.001; Fig. 9D, E). Therefore, we interpreted these results as the more sounds a fish produced

## DISCUSSION

We describe courtship, spawning and aggressive behaviors for *D. dracula*, the most comprehensive description to date of social behavior for any species within the *Danionella* genus of miniature vertebrates. This includes a resident-intruder assay for size-matched and size-mismatched dyads of adult males to quantitatively characterize two prominent display behaviors, sound production and lower jaw extension, that individual males produce during aggressive interactions. We report that *D. dracula* males produce sounds during agonistic, but not courtship interactions. Females are not sonic in either of these contexts or in dyad interactions with males, consistent with their apparent lack of a sound producing apparatus (Britz and Conway, 2016).

The results support the hypothesis that residency results in increased total sound production. Relatively larger fish irrespective of resident or intruder status in size-mismatched assays also exhibit increased total sound production, directed towards the smaller fish in the dyad. The results do not support the hypothesis that jaw extension is a necessary condition for sound production; there is strong support for their being independent displays. We find, however, that resident males in both contexts exhibit increased jaw extension and that time periods of highest sound production and jaw extension often overlapped. Relatively larger males in size-mismatched contexts also exhibit increased aggressive behavior, with significantly increased sound production and jaw extension compared to relatively smaller fish. We also show that when sound production is high, males are making more multi-pulse bursts. In aggregate, the results show that both acoustic and postural displays can serve as robust behavioral benchmarks for comparative studies of *Danionella* species identifying neural, genomic and developmental mechanisms of social behavior evolution within a new model clade (cf. Jourjine and Hoekstra, 2021).

### *Danionella* courtship and spawning

Courtship and spawning behaviors of *D*. *dracula* may differ from those of *D. cerebrum*, the only other *Danionella* species for which these behaviors have been so far reported (Schulze et al., 2018). Female *D. cerebrum* are described to be the first to enter nest tubes, whereupon males follow and spawning occurs. In *D. dracula*, the male is the first to venture into the spawning crevices, followed by the female. In addition, *D. cerebrum* nest tubes are transparent and allow light in, whereas *D. dracula* requires nest coverings that create dark spawning crevices. There is also no description of nest site circling in *D. cerebrum*, whereas *D. dracula* resident males swim close to nest sites from where they court females. These possible differences in courtship and spawning behaviors could parallel ones in the natural habitats of the two species. *Danionella cerebrum* is found in southern Myanmar and *D. dracula* in the north, which may differ in abiotic properties as well as vulnerability to predation. These are all behavioral ecological questions that need to be addressed in the future, especially once these sites become more accessible.

### Acoustic signaling

Individual sound pulses are broadband, short in duration and possess a peak frequency close to 2 kHz in all social contexts observed here. The frequency spectrum of these pulses (Fig. 5) is well within the range of hearing reported for other closely related otophysan genera that, like *Danionella*, have a Weberian apparatus for enhanced sound detection (Fay, 1988; Ladich, 2000; also see Braun and Grande, 2008; Britz and Conway 2016; Conway, Kubicek and Britz, 2021). For some teleost species, dominant frequency correlates with body size as well as with winning in dyadic contests (see Conti et al., 2015 for overview). This is not the case for *D. dracula*, at least under the conditions tested here.

For both size-matched and size-mismatched dyads, males predominantly make single pulse bursts, with repetitive multi-pulse bursts generated during periods of escalated sound production. Bursts of two or more pulses might signal aggressive escalation compared to single pulse bursts, which were more frequently used by all primary aggressor males. *Danionella dracula*’s single pulses are reminiscent of the agonistic “pops” of the bicolor damselfish, *Eupomacentrus partitus* (Myrberg, 1972). Many fish species produce multi-pulse sounds, commonly referred to as “grunts”, during reproductive and aggressive interactions (Lobel, Kaatz and Rice, 2010, Amorim and Almada, 2005; Ladich and Myrberg, 2006; Myrberg and Lugli, 2006; McIver et al., 2014; Lobel et al., 2021). Several damselfish species, including *E*. *partitus*, make multi-pulse “chirps” during reproductive interactions (Myrberg, Spanier and Ha, 1978).

Although *D. dracula* males vary in multi-pulse burst usage and the overall total number of sound pulses produced during male-male dyadic interactions, there is no clear relationship to body size. Future studies of *Danionella* building on the foundation set out here should examine these acoustic characteristics in the context of nest-holding in community tanks to determine if nest-holding males differ in these characteristics compared to “loser” males, similar to other species.

The *Danionella* genus contains multiple species for comparative studies of both sound production and hearing (Britz, Conway and Rüber, 2021). So far, sound production has only been described in one other *Danionella* species, *D. cerebrum* (Schulze et al., 2018). Male *D. dracula* and *D. cerebrum* both make very short broadband, sharp onset pulses around 2 ms in duration. They also exhibit variation in how frequently they sequence bursts during male-male interactions. The sounds of both species can be grouped into bursts separated by a characteristic IPI, close to 35 ms in *D. dracula* and 8 and 17 ms in *D. cerebrum* (Schulze et al., 2018). Beyond the difference in IPI duration, there is a large species difference in the duration of individual bursts and burst trains. We find that *D. dracula* primarily make single pulse signals compared to multi-pulse bursts typically of 2-3 and rarely 4-6 pulses, at least under the conditions tested here. By contrast, *D. cerebrum* have multi-pulse bursts lasting close to 1 s and repetitive bursts lasting on the order of minutes (Schulze *et al.*, 2018). For both species, it would be of interest to compare behaviors that might be coordinated with sound production. The temporal relationship between sound production and lunging is not described for *D. cerebrum* (Schulze et al., 2018) as documented here for *D. dracula*. In addition, *D. cerebrum* lacks the hypertrophied jaw and thus the dramatic jaw extension of *D. dracula*, but perhaps it generates another type of jaw movement during aggressive interactions.

### Jaw extension

Beyond acoustic signaling, animals use other sensory modalities to communicate in conflict scenarios, including visual mechanisms like ornaments and display postures (see Kodric-Brown, Sibly and Brown, 2006; Lappin et al., 2006). As we demonstrate, *D. dracula* males often extend their hypertrophied lower jaw during aggressive interactions. This morphological and behavioral character appears to be unique to this species within the genus. Future studies should test if jaw extension behavior functions as a visual agonistic signal, possibly allowing males to signal contest escalation and assessment information to their competitors. Jaw extension could also act to enhance characteristics of the acoustic signal, akin to the influence of vocal tract and mouth skeleton on the spectral structure and trill rates in songbirds (Podos, Huber and Taft, 2004).

Despite our initial observations that jaw extension was always linked to sound production, we found this to not be the case. The varied temporal separation between the two types of display, especially during periods of low sound production, as well as the independent occurrence of each during male aggressive encounters, indicate these two actions are separate displays. However, the overlap of the two behaviors during periods of heightened activity provides the opportunity to study a possible multimodal signaling repertoire (e.g., see Elias et al., 2003, Amorim et al., 2019) that could maximize robustness of the overall multichannel signal (Ay, Flack and Krakauer, 2007). Males in other sonic species of fish have multimodal signaling repertoires that combine acoustic signals generated by one or more mechanisms (Rice et al., 2022). Sonic signals might also be combined with specific features of other sensory modalities, such as color and ornamentation as well as stereotyped visual dancing/movement displays (Hebets and Uetz, 2000; Elias et al., 2003; Soma and Garamszegi, 2015). Jaw extension by *D. dracula* males could serve as a modifier or amplifier of sound production as well, not contributing its own information but instead augmenting the acoustic signal (Bradbury and Vehrencamp, 2011; Gualla, Cermelli, and Castellano, 2008; Lappin et al., 2006). For a more rigorous investigation of possible temporal coupling/coordination of acoustic and postural displays in *D. dracula* and hence multimodal signaling, the concurrency of sound production and jaw extension should be best re-visited with higher resolution video than used here to examine the timing more precisely between these displays, as done for other sonic species (e.g., Bostwick and Prum, 2003; Fusani et al., 2007).

### Individual Assessment

Resources such as shelters and territories are continually defended by an owner to gain fitness advantages (Conti et al., 2015; Arnott and Elwood, 2007). *Danionella dracula* males swim closely around nest sites that contain spawning crevices. Future studies would benefit from examining if and how males might utilize acoustic communication and other signaling modalities (e.g., vision) to structure dominance hierarchies and determine nest ownership in a community (Conti et al., 2015; Chase et al., 2002, Amorim and Almada, 2005, Arnott and Elwood, 2009, Barata et al., 2007, Myrberg and Riggio, 1985). It would also be essential to determine how nest-holding males might differ from other males in complex acoustic parameters like IPI and multi-pulse burst usage, as well as factors that contribute to fighting potential examined here like nest ownership and body size. Ownership is a factor that can contribute to an overall animal’s motivation, which can often be inferred from competitors’ display rate, or continued agonistic engagement and escalation over resource items, and it can be influenced by internal physiology and perceived resource quality (Bradbury and Vehrencamp, 2011; Arnott and Elwood, 2007; Brown, Chimenti, and Siebert, 2007; Lindström, 1992). How does ownership of a nest site in a community setting affect fighting strategy and escalation, as could be indicated by display rate and sonic characteristics in *D. dracula?* And is it the largest male in the community that holds this nest site? Body size is often a determinant of winning an escalated fight in most animal species, especially those with high variation in body size (Andersson, 1994). A display posture can advantageously reveal body size, and larger animals can produce or bear relatively larger ornaments (Bro-Jørgensen, 2009). We see a size effect in *D. dracula* where the larger male most often (7/8 dyads) is the primary aggressor in size-mismatched dyads. This size asymmetry effect in *D. dracula* contests is reminiscent of Assessor-like strategies (Parker, 1974; Arnott and Elwood, 2009) and resembles other species which can assess differences in body size (see Introduction). In size-matched dyads however, we found a distinct resident effect in the absence of size differences between males, indicating the possible presence of a strategy in *D. dracula* contests where residents are predicted to escalate aggression compared to intruders (Maynard Smith, 1979), and so there may be possible tradeoffs between residency and size asymmetry in *D. dracula* fighting strategies (Hack, Thompson and Fernandes, 2010; Hofmann and Schildberger, 2001; Jennions and Backwell, 1996, Jackson and Cooper, 1991).

### Concluding Comments

We demonstrate that *D. dracula* is a sonic species and characterize the sounds produced by individual males in dyadic assays for size-matched and size-mismatched contexts, uncovering initial residency and body size effects on aggressive behaviors in this species. Future physiological investigations of auditory sensitivity (see Bass and McKibben, 2003), as well as whole-brain neuroimaging of *Danionella* (Schulze et al., 2018; Chow et al., 2020), can determine the components of natural acoustic and visual/postural signals that are attended to by both males and females. The results presented here are a basis for subsequent behavioral studies determining how *Danionella* species in general may assess each other’s fighting ability via multiple sensory modalities, such as audition and vision for observing sonic and postural (jaw extension) behaviors, respectively. The temporal relationship of sound production to other displays such as jaw extension could also allow researchers to determine how such actions may augment the information conveyed through sound production, perhaps acting as a tactical threat or amplifier (see Bradbury and Vehrencamp, 2011). Last, our experiments provide a foundation on which to test established social behavioral models of conflict resolution such as fighting strategy models (see Bradbury and Vehrencamp, 2011), which altogether would make a comprehensive framework on which to study the neural and genetic drivers of social behaviors in *Danionella* species.

## ACKNOWEDGEMENTS

We sincerely thank Kevin Conway for his early and very generous guidance in setting up a breeding tank of *D. dracula*, and Michael Sheehan for many helpful comments on the manuscript. We also thank the Cornell Statistical Consulting Unit for discussions on data analysis. We also thank Melissa Hoffman for animal care and Margaret Marchaterre for logistical support.

## FOOTNOTES

### Author Contributions

Conceptualization: R.L.T., A.H.B., H.K.R.; Methodology: R.L.T., A.H.B, S.C.C., J.T.P, E.R.S., H.K.R; Validation: R.L.T.; Statistical analyses: R.L.T., E.R.S., H.K.R; Data aggregation: R.L.T., Z.G., S.C.C., J.T.P., H.K., W.F; Data curation: R.L.T.; Writing - initial draft: R.L.T., A.H.B.; Revision: R.L.T., A.H.B, Z.G., S.C.C., J.T.P, E.R.S., H.K.R; Visualization: R.L.T.; Resources, Supervision, Project administration, Funding acquisition: A.H.B. All authors approved the final version of the manuscript.

