## Supplementary Figures for "Acoustic and postural displays in a miniature and transparent teleost fish, *Danionella dracula*"

SUPPLEMENTARY FIGURES AND MOVIES

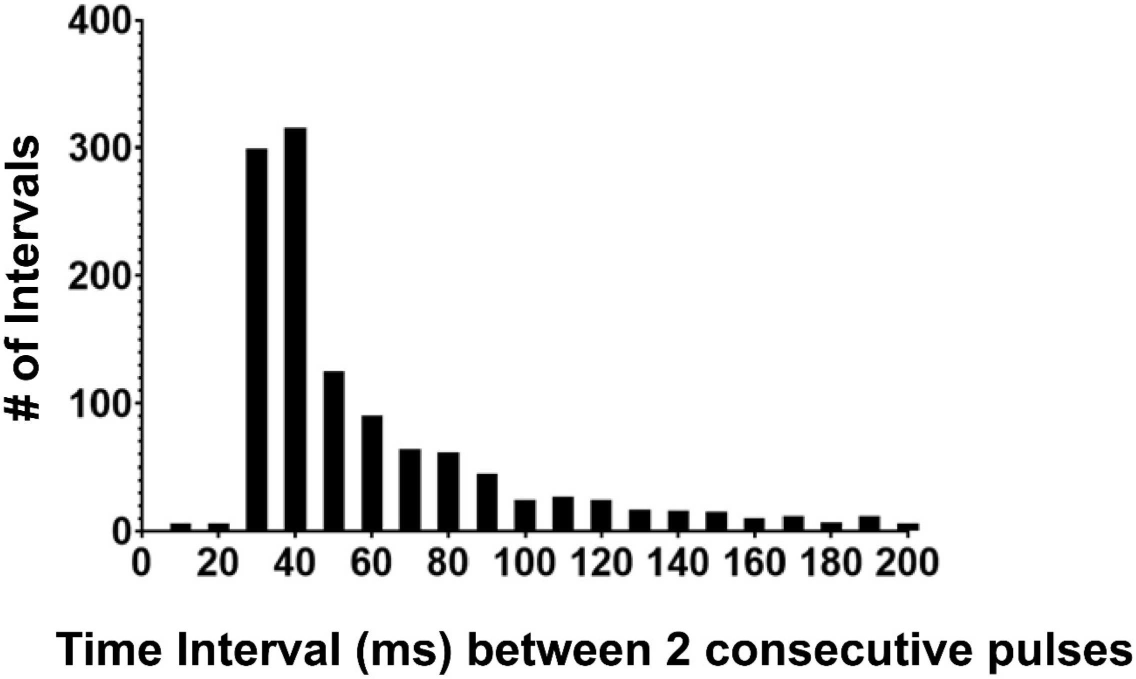


Figure S1: Frequency distribution of intervals (ms) measured between 2 consecutive individual pulses (all intervals from both size-matched and size-mismatched contexts). The mode and peak is 34 ms. Data is visualized through 200ms for ease of visibility.

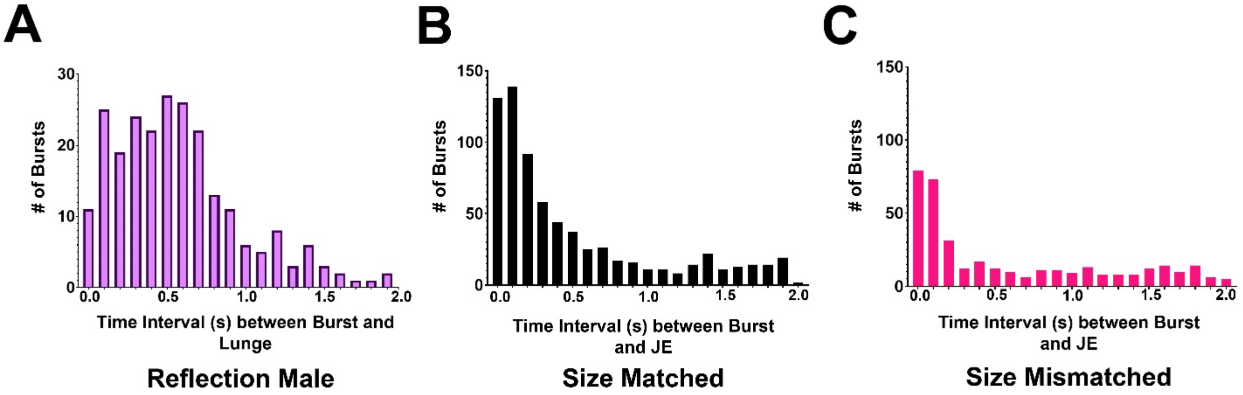


Figure S2: Frequency distribution of intervals (ms) measured between 2 consecutive individual behavioral events. A. Frequency distribution of intervals (s) measured between a burst of sound and a lunging event. B. Frequency distribution of intervals (s) measured between a burst of sound and an extension of the lower jaw in high sound producing and high jaw extending males in size-matched contests (n=5). C. Frequency distribution of intervals (s) measured between a burst of sound and an extension of the lower jaw in high sound producing and high jaw extending males (n=3) in size-mismatched contests.

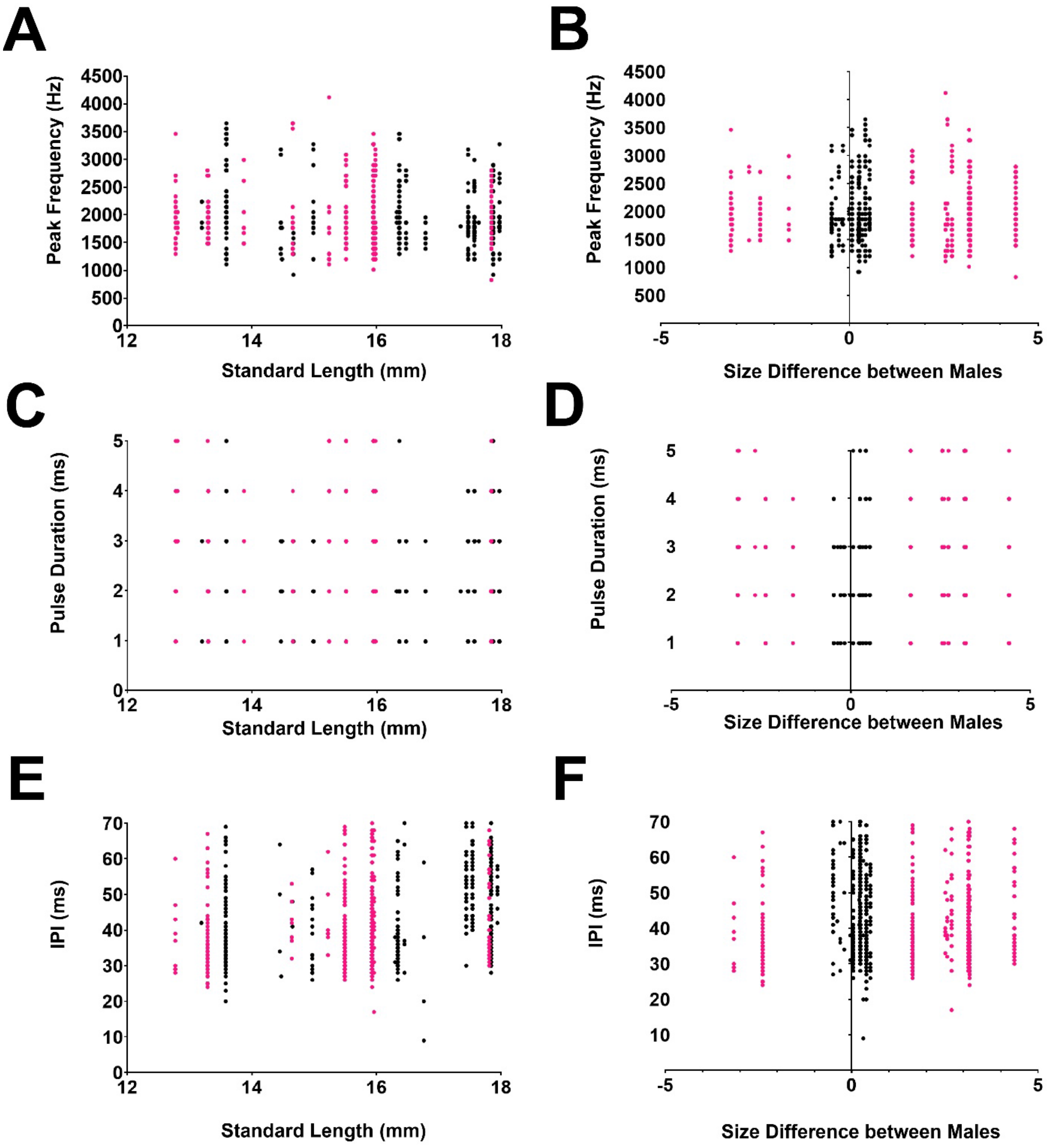


Figure S3: Scatterplots of individual males' standard length (mm) or size difference from competitor (mm) by sound characteristics of peak frequency (Hz), pulse duration (ms) and inter-pulse interval (ms). Black circles indicate measurements from individual males in size-matched contests; Salmon pink circles indicate measurements from individual males in size-mismatched contests. A. Individual male's standard length (mm) plotted by peak frequency (Hz) of individual pulses (n=1259 pulses in size-matched contests; n=1453 pulses in size-mismatched contests). B. Individual male's size difference from competitor (mm) plotted by peak frequency (Hz) of individual pulses (n=1259 pulses in size-matched contests; n=1453 pulses in size-mismatched contests). C. Individual male's standard length (mm) plotted by pulse duration (ms) of individual pulses (n=1259 pulses in size-matched contests; n=1453 pulses in size-mismatched contests). D. Individual male's size difference from competitor (mm) plotted by pulse duration (ms) of individual pulses (n=1259 pulses in size-matched contests; n=1453 pulses in size-mismatched contests). E. Individual male's standard length (mm) plotted by inter-pulse interval (IPI) (ms) of individual pulses (n=422 pulses in size-matched contests; n=456 pulses in size-mismatched contests). F. Individual male's size difference from competitor (mm) plotted by by inter-pulse interval (IPI) (ms) of individual pulses (n=422 pulses in size-matched contests; n=456 pulses in size-mismatched contests).

Table S1. Community Tank Behavioral Events for *D. dracula*

| ***Event*** | **Definition** |
|---|---|
| Lunge at male | Focal male orients head towards and swims rapidly towards male. Trajectory directed at target male but does not pursue as target swims away. |
| Courtship | Focal male swims beneath female and vibrates its head and fins back and forth underneath egg vent. |
| Enter nest | Focal male swims into crevice of nest head-first. |

Table S2. Principal component analysis of multi-pulse burst in size-matched and size-mismatched dyadic contests: Loading Coefficients.

| | Size-matched | Size-mismatched |
|---|---|---|
| **Type of multi-pulse burst** | **PC1** (77%) | **PC1** (65%) |
| Two pulse | 0.96 | 0.97 |
| Three pulse | 0.99 | 0.92 |
| Four pulse | 0.92 | 0.68 |
| Five pulse | 0.86 | 0.59 |
| Six pulse | 0.59 | ---- * |

*Six pulse bursts were never observed in size-mismatched contests

Movie 1: A male fish lunges at its own reflection in the tank wall, producing sounds and exhibiting extension of the lower jaw.

Movie 2: Male swims below female and vibrates his body and head back and forth beneath the female’s egg vent. Male then swims back to nest entry crevice and female swims behind male to same nest entry crevice. Movie zooms in to show male as he swims headfirst into nest entry crevice and female orients head towards crevice before swimming headfirst into crevice.

Movie 3: In a dyadic interaction, a male orients head towards other male and lunges at other male, producing a series of pulses and extending his lower jaw. The other male swims away from the lunging male.
